# Nuclear and mitochondrial phylogeny of *Rossella* (Hexactinellida: Lyssacinosida, Rossellidae): a species and a species flock in the Southern Ocean

**DOI:** 10.1101/037440

**Authors:** Sergio Vargas, Martin Dohrmann, Christian Göecke, Dorte Janussen, Gert Wöerheide

## Abstract

Hexactinellida (glass sponges) are abundant and important components of Antarctic benthic communities. However, the relationships and systematics within the common genus *Rossella* Carter, 1872 (Lyssacinosida: Rossellidae) are unclear and in need of revision. The species content of this genus has changed dramatically over the years depending on the criteria used by the taxonomic authority consulted. *Rossella* was formerly regarded as a putatively monophyletic group distributed in the Southern Ocean and the North Atlantic. However, molecular phylogenetic analyses have shown that *Rossella* is restricted to the Southern Ocean, where it shows a circum-Antarctic and subantarctic distribution. Herein, we provide a molecular phylogenetic analysis of the genus *Rossella*, based on mitochondrial (16S rDNA and COI) and nuclear (28S rDNA) markers. We corroborate the monophyly of *Rossella* and provide evidence supporting the existence of one species, namely *Rossella antarctica* Carter, 1872 and a species flock including specimens determined as *Rossella racovitzae* Topsent, 1901, *Rossella nuda* Topsent, 1901, *Rossella fibulata* Schulze & Kirkpatrick, 1910, and *Rossella levis* (Kirkpatrick, 1907).

## 1 Introduction

Glass sponges (class Hexactinellida) are key components of Antarctic suspension-feeder communities (Arnaud et al., 1998; Gutt, 2007). Antarctic hexactinellids, particularly species of the genera *Rossella* and *Anoxycalyx (Scolymastra)*, can reach remarkable size, biomass and abundance (Barthel and Tendal, 1994; Janussen and Tendal, 2007; McClintock et al., 2005). At some localities, *Rossella* species dominate the seafloor (Fig. 1), and thus increase its spatial heterogeneity (Gutt and Starmans, 1998; Janussen and Reiswig, 2009; Starmans et al., 1999; Dayton et al., 2013; Fillinger et al. 2013), structure benthic communities (Barthel, 1992a, b), and locally play a major role in silicon cycling (Gatti, 2002; Gutt et al. 2013). Large *Rossella* specimens can harbour a diverse community of invertebrates and juvenile stages of many other organisms, and serve as substratum for various taxa of other sessile invertebrates (epibionts) (Barthel, 1997; Gutt and Schickan, 1998; Kunzmann, 1996; Kersken et al., 2014).

**Figure 1:**
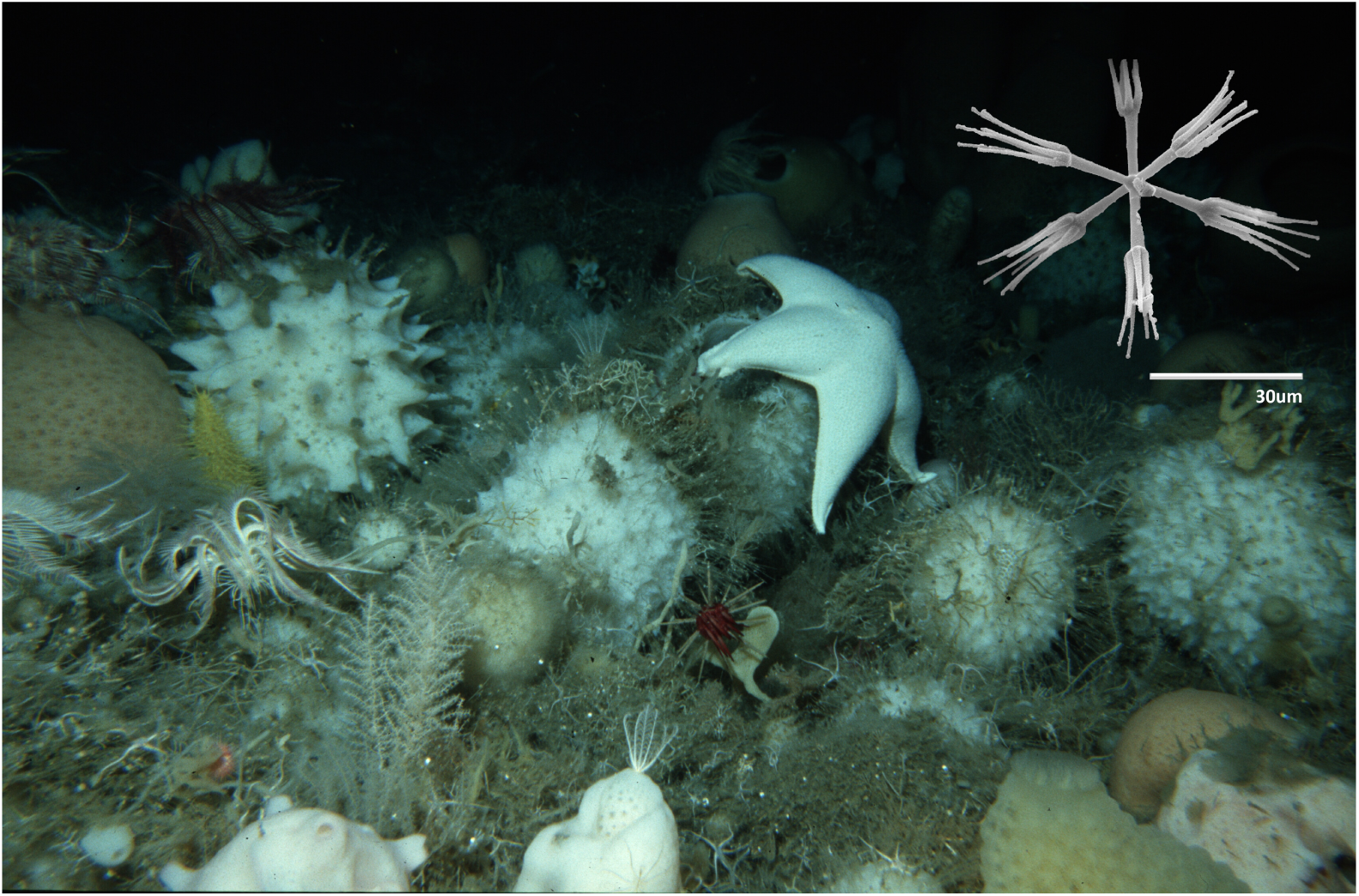
Seafloor dominated by *Rossella* spp. in the Weddell Sea, Antarctica (latitude: -70.8717, longitude: -10.5233; depth=254m); Photo: Gutt, J. and Starmans, A. (2004): Sea-bed photographs (benthos) along ROV profile PS56/127-1. doi:10.1594/PANGAEA.198695. Inset: calycocome of *Rossella antarctica*; Photo: C. Göcke.

From a morphological perspective, *Rossella* is characterized by the presence of calycocomes (Fig. 1) and its typical microhexasters (Tabachnick, 2002), although similar spicules also occur in a few other genera (see Dohrmann et al., 2012). In contrast to the relatively stable genus-level systematics, intra-generic relationships remain unclear and most *Rossella* species still require a revised and clear morphological delineation (Barthel and Tendal, 1994, Göcke and Janussen, 2013). As a result, the number of species recognized for the genus has varied in the past, ranging from two to 21 species depending on the taxonomic authority (e.g. Burton, 1929; Barthel and Tendal, 1994; Koltun, 1976; see also van Soest, et al. 2015). The great variation of the number of recognized species is, to some extent, not surprising. With the sole exception of *Rossella antarctica*, all remaining species lack clear morphological apomorphies (cf. Barthel and Tendal, 1994), or their diagnostic characters are weak (e.g. external morphology, shape and size of dermal megascleres). In addition, the majority of characters used for species delimitation in the genus are continuous, making differences between species mainly gradual and subject to diverse interpretations. Calycocome sizes, for instance, tend to overlap between species, as does the size of other taxonomically important spicules (Barthel and Tendal, 1994; Göcke and Janussen, 2013). External body shape is variable even within species (Tabachnick, 2002). Finally, the lack of appropriate sampling has, to some extent, hampered the systematic evaluation of the variability of the main characters used for distinguishing different species.

Clarifying the systematic relationships within the genus *Rossella* is an important task of potential benefit to other areas of Antarctic research. *Rossella* species are structurally important in Antarctica (see above), and their distribution, as that of many other Antarctic sponges, is thought to be circum-antarctic (Janussen and Reiswig, 2009; Sara et al., 1992). The role that different *Rossella* species play in structuring Antarctic communities, as well as whether some or all species are, indeed, a circumpolar cohesive unit, strongly depends upon the clear delineation of those species. Here, we provide a phylogenetic analysis of the genus *Rossella* based on mitochondrial (16S rDNA, COI) and nuclear (28S rDNA) markers and including 5 of the 8 species currently recognized as valid by Barthel and Tendal (1994). We aim to test different morphology-based taxonomic arrangements that have been proposed for the genus throughout its taxo-nomic history and attempt to reconcile the current morphology-based classification of *Rossella* spp. from the Antarctic Weddell Sea with the molecular results obtained here to further clarify the evolution of this important SO taxon.

## 2 Materials and Methods

### 2.1 Specimens and laboratory procedures

Specimens (Table 4) were collected by trawling during the German ANT XXIII/8 (2006/2007) and ANT XXIV/2-SYSTCO Expedition (2007/2008) to the Weddell Sea (Atlantic sector of West Antarctica), photographed and fixed in 96% ethanol. All specimens are deposited in the Senckenberg Naturmuseum (Frankfurt a.M., Germany) and were catalogued in the online database SESAM. Sponges were determined to species level using standard procedures (e.g. Janussen et al., 2004) and pertinent literature. DNA was extracted from small pieces of tissue with the NucleoSpin DNA tissue extraction kit (Macherey-Nagel) following the manufacturer’s protocol.

**Table 4:**
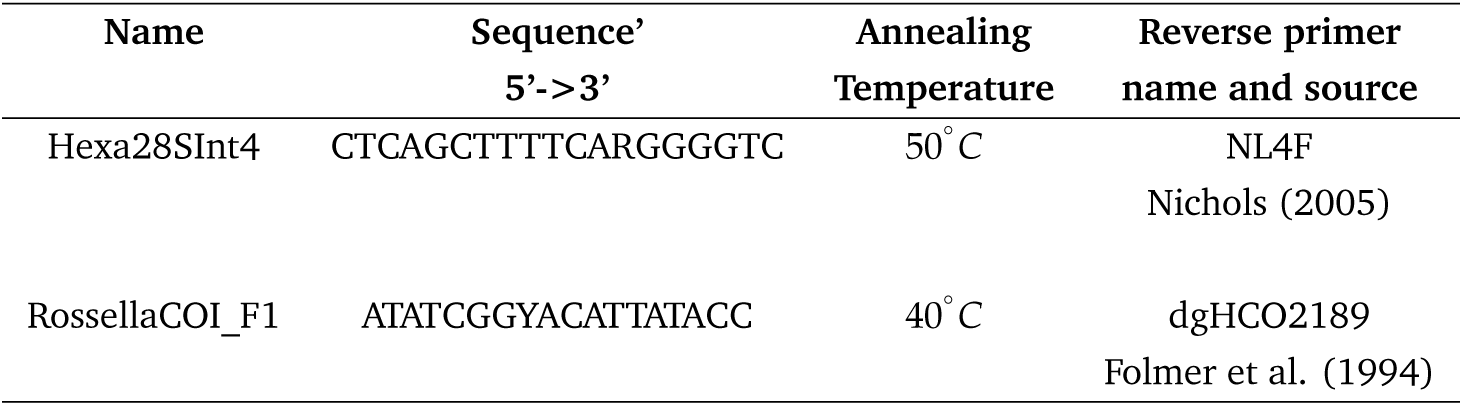
Primers and annealing temperatures used for amplification of 28S rDNA and COI.

Three different molecular markers —partial 28S rDNA (ca. 1.2kb), partial 16S rDNA (ca. 0.5kb), and the standard barcoding fragment (Folmer et al. 1994) of COI (ca. 0.6kb)— were amplified using 12.5 *μ*l reaction volumes of GoTaq (Promega) supplemented with BSA. Three-step PCR protocols, including an initial denaturation step of 94 °C 3 min, 35 to 40 cycles of 94 °C 30 s, 50 °C/40 °C 30 s, 72 °C 60 s, and a final extension of 5 min at 72 °C were used for all markers (see the supplementary materials for details on the annealing temperature for each primer). For 28S rDNA and COI we designed *Rossella-*specific primers (see Suppl. Materials) to avoid co-amplification of non-target organisms; 16S rDNA primers were as in Dohrmann et al. 2008. PCR products were cleaned by standard ammonium acetate-ethanol precipitation or ExoSap-IT (Affymetrix) enzymatic PCR clean-up and sequenced in both directions using the same primers used for PCR and the BigDye Terminator 3.1 chemistry (Applied Biosystems). Sequencing reactions were precipitated with sodium acetate-ethanol and subsequently analyzed on an ABI 3700 Genetic Analyzer at the Sequencing Service of the Department of Biology, LMU München. Trace files were assembled in CodonCode Aligner (CodonCode Corporation); hexactinellid origin of all obtained sequences was verified using NCBI BLAST (Johnson et al., 2008). Sequences are deposited at EMBL under accession numbers HE80191 to HE80223.

### 2.2 Outgroup choice and sequence alignment

New sequences were manually aligned in SeaView 4 (Gouy et al., 2010) to published alignments (Dohrmann et al., 2012b). However, we restricted the taxon set to representatives of the families Leucopsacidae and Rossellidae, as well as *Clathrochone clathroclada* (Lyssacinosida incertae sedis). Leucopsacidae and *C. clathroclada* have been shown to be successive sister groups to Rossellidae (Dohrmann et al. 2012b), and were therefore used as outgroups. Alignments were concatenated into a supermatrix and ambiguously alignable regions removed. The final alignment is 1.2 kb long and is available at https://bitbucket.org/sevragorgia/rossella.

### 2.3 Phylogenetic analysis

Using the concatenated alignment, we inferred both Maximum likelihood (ML) and Bayesian phylogenies with RAxML 7.2.8 (Stamatakis, 2006) and PHASE 2.0, respectively. The GTR model of nucleotide substitution (Tavaré, 1986) was used for 16S rRNA, COI as well as for 28S rRNA single-stranded regions (loops). Among-site rate variation was modelled using a discrete approximation of a gamma distribution with 4 categories (+G; Yang, 1994, 1996). For the stem regions (paired sites) of the 28S rRNA we used the S16 and S7A models of sequence evolution (Savill et al., 2001) for the ML analysis. We searched for the ML tree using 20 independent tree-search replicates and assessed branch support with 1000 bootstrap pseudo-replicates (Felsenstein, 1985), using the “rapid bootstrap” algorithm described by Stamatakis et al. (2008). In the Bayesian analysis, two independent Markov Chain Monte Carlo (MCMC) chains were run for 10,000,000 generations after a burn-in of 250,000 generations, sampling every 100 generations. Model specifications for the Bayesian analysis were the same as for the ML analysis (i.e. GTR+G for 16SrDNA and COI); however we only used the S7A model for the 28S rRNA stem regions because it was difficult to achieve chain convergence using the S16 model.

### 2.4 Partition addition bootstrap and alternative lineage attachment analysis

To assess the influence of the individual partitions or combinations thereof on the ML topology inferred from the concatenated data matrix (i.e. the total evidence ML tree), we performed ML bootstrap analyses (1000 pseudoreplicates) for each individual marker and for all combinations of two markers using RAxML 7.2.8. For all these analyses, we used the same model settings as in the total evidence analysis for the corresponding partition (e.g. GTR+G for 16S rDNA and S16 for 28S rDNA stems). After each analysis, we determined the partition-specific bootstrap support (sensu Struck et al., 2006) for the branches present in the total evidence tree using consensus from the phyutility package (Smith and Dunn, 2008). We also assessed alternative branching positions of different *Rossella* species using linmove from the phyutility package. In brief, linmove screens a set of phylogenetic trees and reports the frequency with which alternative placements of a branch occur in that set. The analysis facilitates the visualization of alternative branching positions of a lineage showing low bootstrap support values, which allows to determine whether poorly supported branches have only a few attachment points occurring with high frequency or branch off at several multiple positions with low frequency.

### 2.5 Testing hypotheses of relationships within *Rossella*

Different taxonomic arrangements proposed for *Rossella* can be translated into specific phylogenetic hypotheses and evaluated with available statistical tests (e.g., Goldman et al., 2000; Huelsenbeck, 1997; Huelsenbeck and Crandall, 1997; Whelan et al., 2001). We used the AU test implemented in CONSEL (Shimodaira and Hasegawa, 2001; Shimodaira, 2002) with site-wise log-likelihood values obtained from RAxML 8.2.4 to test the monophyly of *R. nuda* and *R. racovitzae*, the two species for which more that one specimen was available. Briefly, ML analyses constrained to enforce the monophyly of *R. nuda, R. racovitze* or of these two species were performed in RAxML and the best constrained ML trees were compared against the best unconstrained phylogeny.

## 3 Results

### 3.1 Total evidence phylogenetic analysis

We recovered a phylogenetic tree congruent with published analyses of the class Hexactinellida (Dohrmann et al., 2008, 2009, 2012a,b). Bayesian and ML analyses recovered generally similar trees, but the Bayesian phylogeny did not include a clade of *R. fibulata* + *R. racovitzae*, which was present in the ML phylogeny with moderate support. Both independent MCMC runs of the Bayesian analysis converged to the same consensus topology. The ML topology was not sensitive to model choice for the 28S rDNA stem sites, as analyses using S7A and S16 resulted in the same phylogeny.

Our phylogenetic analyses (Fig. 2) recovered a well supported clade comprising all *Rossella* spp. which nested deeply within the family Rossellidae. *Rossella antarctica* specimens formed a highly supported clade in both Bayesian and ML analyses. Specimens belonging to other morphologically defined *Rossella* species formed a large clade hereafter named the *Rossella racovitzae* clade. Within this clade, other morphologically defined *Rossella* species were not recovered as monophyletic, but were polyphyletic in both the ML and Bayesian tree. Support values within the *R. racovitzae* clade were generally low (< 50 %), with only some branches showing moderate (50 - 80 %) bootstrap support in the ML analysis. In contrast, the Bayesian analysis assigned high posterior probabilities (PP > 0.95) to most branches within this clade.

**Figure 2:**
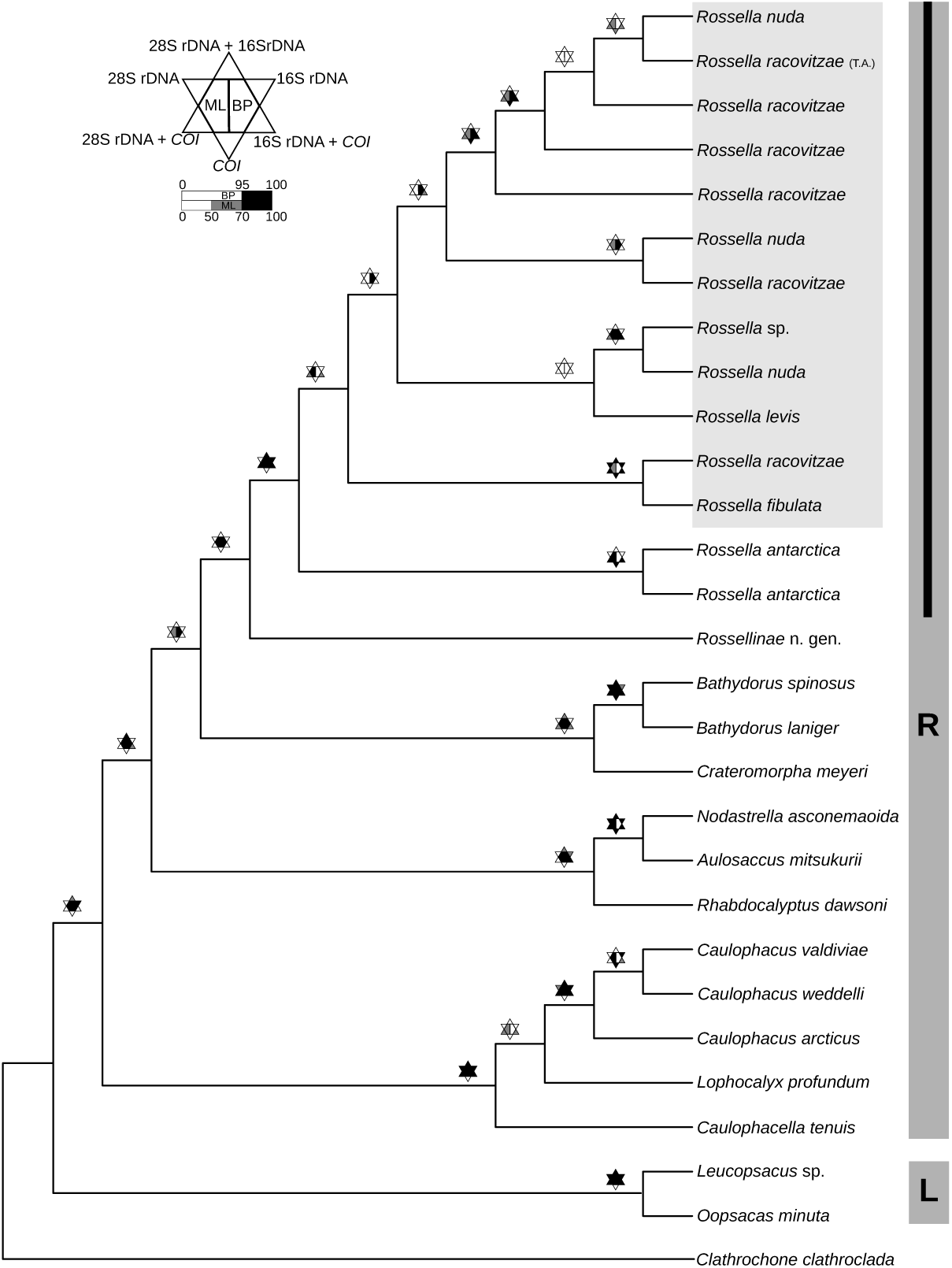
Phyloenetic relationships (cladogram) of Southern Ocean *Rossella*. The tree corresponds to the total evidence maximum likelihood topology. The vertices of the stars above the branches show the bootstrap value obtained for a given branch when using a single partition or a combination of partitions (see inset). The centers of the stars show, on the left, the bootstrap value of the maximum likelihood total evidence analysis, and on the right, the posterior probability obtained for the branch in the Bayesian analysis. Dark gray bars on the right annotate the family Rossellidae (R) and Leucopsacidae (L); Clathrochone is currently *incertae sedis* in Lyssacinosida. Within Rossellidae, *Rossella* is indicated with a black bar and *Rossella racovitzae sensu lato* highlighted in ligth gray. Information about other specimens included in the analysis can be found in Dohrmann et al. (2008, 2009). Both topologies, ML and Bayesian, as well as partition specific trees with branch-lengths and support values are provided in the Suppl. Materials.

### 3.2 Partition addition bootstrap analysis and lineage movement

Bootstrap values assigned to the branches of the total evidence ML tree varied between different partitions or combinations thereof (Fig. 2). In general, bootstrap support increased when more data were added to the analysis. However, there was conflict between partitions in some specific cases. For instance, the *R. antarctica* clade was not supported by 16S rDNA sequences alone but received high bootstrap support from the COI partition. When the two markers were combined, bootstrap support was only moderate (50 - 80%) in contrast to the high support (> 80%) assigned to this clade in the total evidence analysis. Within the *R. racovitzae* clade, support was low when single partitions or combinations of two partitions were used for the analysis, and was only moderate in the total evidence phylogeny. Lineage movement analysis revealed that morphospecies included in the *R. racovitzae* clade were not monophyletic in any of the bootstrap pseudo-replicates, invariably forming clades with specimens belonging to different morphospecies.

### 3.3 Hypothesis testing

All constrained phylogenetic hypotheses explored in this study were found to be significantly worse (p < 0.001) than the unconstrained ML tree (Table 2). The decay in the likelihood values of the constrained ML phylogenies was highly related to the number of constraints. Trees constrained to make single species monophyletic (e.g. *R. nuda* or *R. racovitzae)* showed higher log likelihood values than trees constrained to make all species monophyletic (e.g. *R. nuda* and *R. racovitzae* monophyletic). These results were insensitive to model selection, leading to identical conclusions when either the S16 or the S7A model was applied to 28S rDNA stems.

**Table 1:** Collection details of sequenced specimens.

| Species | Expedition/Station/Voucher number | Coordinates (Latitude;Longitude) | Depth (meters) |
| --- | --- | --- | --- |
| *Rossella racovitzae* | Systco/48-1/SMF11729 | 70° 23.94' S; 8° 19.14' W | 602.1 |
| *Rossella racovitzae* | Systco/48-1/SMF11733 | 70° 23.94' S; 8° 19.14' W | 602.1 |
| *Rossella racovitzae* | Systco/48-1/SMF11736 | 70° 23.94' S; 8° 19.14' W | 602.1 |
| *Rossella nuda* | Systco/48-1/SMF11715 | 70° 23.94' S; 8° 19.14' W | 602.1 |
| *Rossella levis* | Systco/48-1/SMF11728 | 70° 23.94' S; 8° 19.14' W | 602.1 |
| *Rossella fibulata* | Systco/48-1/SMF11732 | 70° 23.94' S; 8° 19.14' W | 602.1 |
| *Rossella antarctica* | Systco/48-1/SMF11734 | 70° 23.94' S; 8° 19.14' W | 602.1 |
| *Rossella antarctica* | Systco/48-1/SMF11735 | 70° 23.94' S; 8° 19.14' W | 602.1 |
| *Rossella racovitzae* | ANTXIII-8/697-1/SMF11731 | 63° 15.38' S; 59° 3.94' W | 143.7 |
| *Rossella nuda* | ANTXIII-8/700-4/SMF11730 | 65° 56.08' S; 60° 20.28' W | 211.1 |
| *Rossella* sp. | STN54AEV393GD4075 (SVR68) | ? | ? |

**Table 2:** Constrained phylogenetic hypotheses tested using the AU-test.

| **Monophyly**<br>**constraint** | **Best**<br>**log likelihood** | **AU-test**<br>**AU-test** |
| --- | --- | --- |
| No constraint | -6178.947402 | N.A. |
| *R. nuda* | -6229.382186 | $p < 0.001$ |
| *R. racovitzae* | -6206.034912 | $p = 0.024$ |
| *R. racovitzae*+*R. nuda* | -6245.485101 | $p < 0.001$ |

## 4 Discussion

The last 50 years of taxonomic history have seen genus *Rossella* expanding from two species, *R. antarctica* and *R. racovitzae* (Rossella-concept of Koltun, 1976), to eight species (Rossella-concept of Barthel and Tendal, 1994) to 19 species currently accepted as valid in the World Porifera Database (van Soest et al. 2015). All six species resurrected by Barthel and Tendal (1994) were included in the broad and ‘highly polymorphic’ *R. racovitzae* by Koltun (1976). Here, we have sequenced two mitochondrial markers and one nuclear marker in an attempt to clarify the systematics of *Rossella* using an independent set of characters not used by previous authors. Our results reveal that genus *Rossella* divides into two main clades corresponding to Koltun’s *Rossella* species: a well supported *R. antarctica* clade was recovered as sister to a moderately supported group of specimens assigned to various nominal species and here referred to as the *R. racovitzae* clade. Both clades have clear diagnostic molecular characters in their COI sequences, i.e., molecular synapomorphies (Fig. 3). The morphology-based taxonomy of the species included within the *R. racovitzae* clade is not straightforward. Most of its species lack clear apomorphic characters and many of the characters used for species delimitation inside this clade overlap or are prone to authoritative (subjective) interpretation (Table 3). In addition, broad morphological variation in both external and spicule morphology is found in the *R. racovitzae* clade (Barthel and Tendal, 1994; Göcke and Janussen, 2013). Yet, the *R. racovitzae* clade can be roughly subdivided into several groups that display morphological cohesiveness and to some extent correspond to the described species included in this clade (Göcke and Janussen, 2013). In contrast, *R. antarctica* can be readily identified and can be clearly distinguished from all other *Rossella* species based on morphology as well as their clear diagnostic molecular characters; in contrast, no diagnostic molecular characters were found for all the other species within the *R. racovitzae* clade in the standard barcoding partition.

**Table 3:**
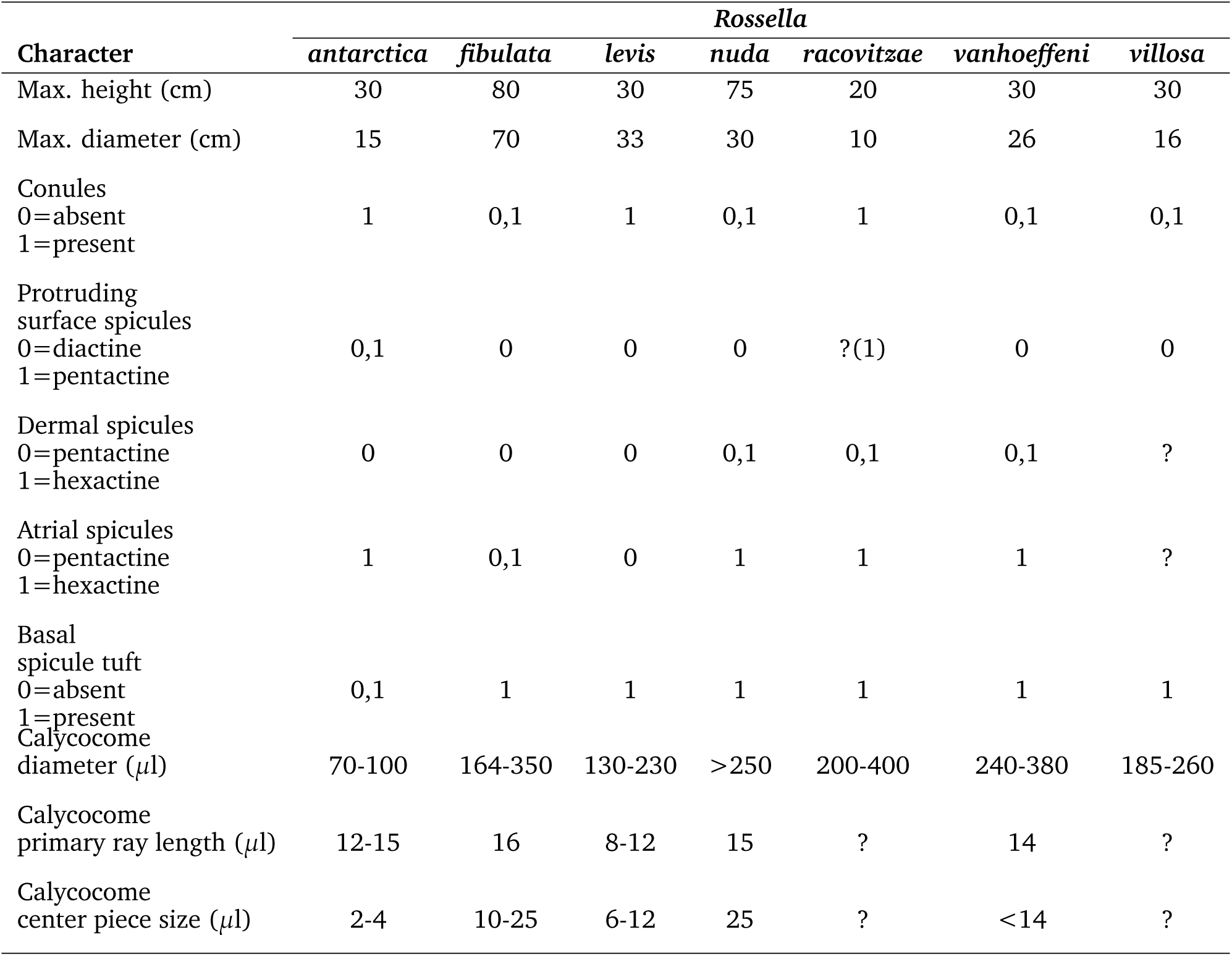
Selected morphological characters used for the taxonomy of the SO *Rossella* after Barthel and Tendal (1994).

**Figure 3:**
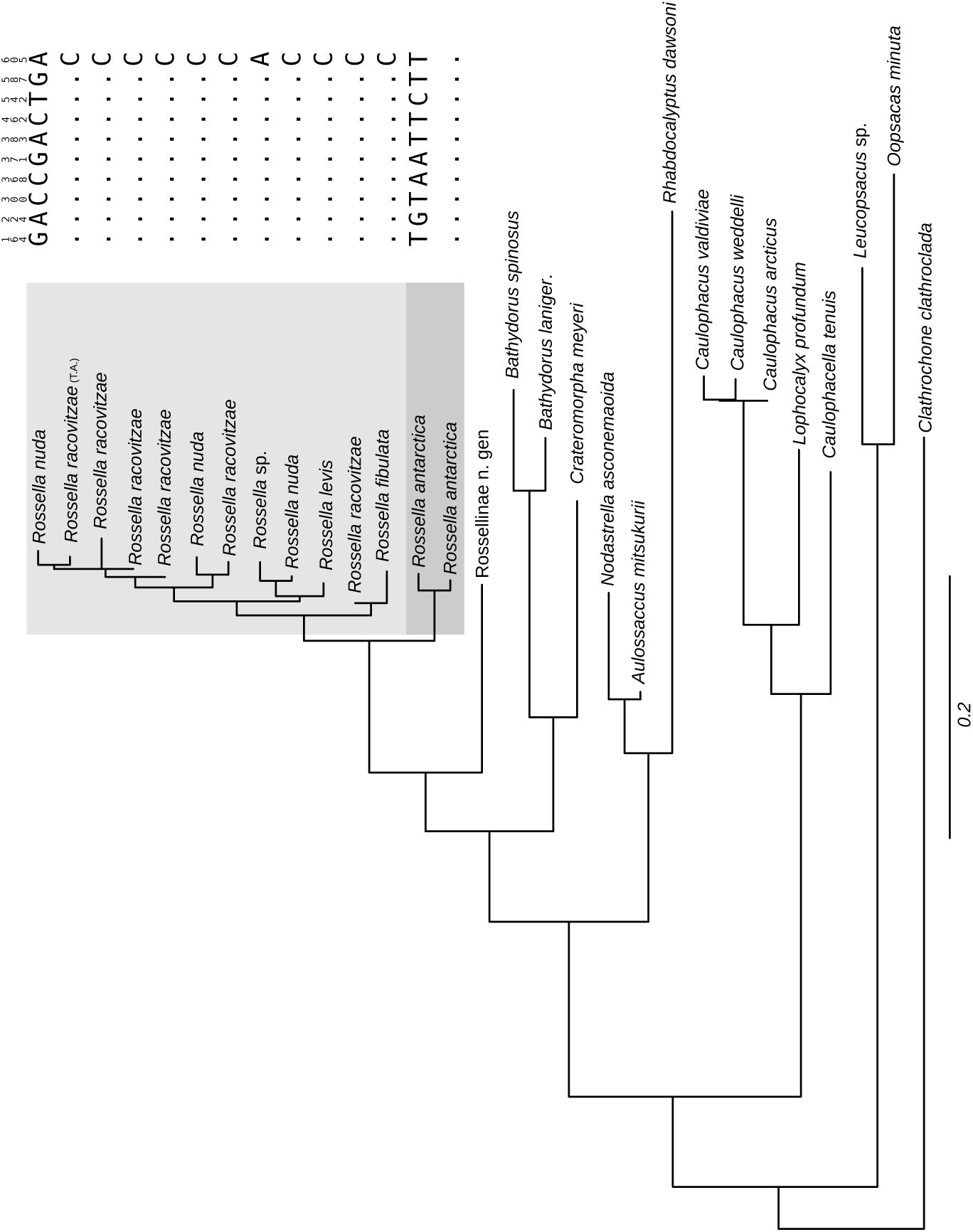
Maximum likelihood phylogram of Southern Ocean *Rossella* based on the total evidence data matrix. For the bootstrap values of the nodes see Figure 1. Highlighted are the two main *Rossella* s.s. clades obtained with their corresponding COI diagnostic characters: consensus sequence on top of the alignment, positions identical to the consensus represented with dots. Scale bar, expected number of substitutions per site.

The analysis of the alternative branching positions of specimens included within the *R. racovitzae* clade revealed that specimens morphologically assigned to the same nominal species were not monophyletic in any bootstrap tree of the total evidence ML analysis nor in the Bayesian tree; the AU tests of monophyly applied to some phylogenetic hypotheses constrained to group different *Rossella* species together also rejected the monophyly of the species tested. Species are expected to be poly- or paraphyletic after or during speciation. Therefore, the non-monophyly within the *Rossella racovitzae* clade in the molecular phylogeny presented here could reflect a recent or ongoing speciation process in the genus. Consequently, we propose that the *R. racovitzae* clade, in contrast to the well defined *R. antarctica*, is a species flock. Species flocks are monophyletic, diverse (morphologically, ecologically, and taxonomically) assemblages of closely related species which evolved rapidly within an area where they are endemic and ecologically dominant (Lecointre et al., 2013). The *R. racovitzae* species flock includes 4 out of 5 species here sampled, these species are endemic to Antarctica, are morphologically diverse and appear to have evolved rapidly as judged by their poly- or paraphyletic status (observed here) and their biogeographic history. Molecular divergence time estimations resulted in a mean age of 40±20 Ma for crown-group *Rossella* (Dohrmann et al. 2013). This age accords well with the opening of the Drake Passage ( 30 Ma), a geological event that resulted in the final isolation of Antarctica (Lawver and Gahagan, 2003) and could have caused the diversification of *Rossella* in this region. We consider this “Rossella-concept", including one clear species and a species-flock, to currently best reconcile all available evidence (i.e. morphological and molecular) and to provide an evolutionary framework to interpret the high levels of variation within the *R. racovitzae* clade without requiring the synonimization of most *Rossella* species in a “highly polymorphic” *Rossella* species (i.e. *R. racovitzae* s.l.).

From a biogeographic perspective, the circum-Antarctic cohesiveness of both *R. antarctica* and members of the *R. racovitzae* clade remains to be tested. In this study, the only specimen of *R. racovitzae* from east Antarctica (collected in Terra Adelie) included in the analysis was sister to specimens from the Weddell Sea. However, any conclusion about the biogeography of *Rossella* species in the SO derived from our current dataset seems premature given the restricted geographic coverage of our sample.

Finally, we would like to highlight the difficulties in getting access to fresh material of all valid species of *Rossella* for molecular phylogenetic analyses. *Rossella* is well known for its abundance in Antarctica, however most specimens collected belong to *R. racovitzae* and specimens beloging other species are collected less frequently. *R. levis* was only collected 3-5 times in 4 expeditions to the Antarctica by DJ and something similar occurs to *R. fibulata* (cf. Göcke and Janussen, 2013). *R. racovitzae* and *R. nuda* are more often collected and more material is generally available from these species. Two other species *R. vanhoeffeni* and *R. villosa* have not been collected after years of field work in the Weddell Sea. This difficulties, somewhat normal in the deep-sea, but paradoxical given the reported abundance of *Rossella* spp. in Antarctic waters, hamper the thorough testing of the monophyly of most species in the genus. We provide here the first molecular study of the phylogenetic relationship within this important sponge genus in Antarctica and pursue to reconcile the morphological and molecular evidence given the available material and to open new avenues for future work to further clarify the phylogeny of *Rossella*.

## 5 Conclusion

We have obtained a phylogeny of the genus *Rossella* corroborating its monophyly and showing the existence of two clades corresponding to the well-defined species *R. antarctica* and a diverse assemblage of species, here considered a species flock and termed *R. racovitzae* flock. Future sampling and the use of genome-wide molecular markers will certainly contribute to expand our understanding of these important Antarctic species, in particular about the circumpolar distribution of *Rossella*, the causes of the high morphological diversity and the relationships within the *R. racovitzae* species flock.

## Acknowledgements

We thank Annamarie Gabrenya and Astrid Schuster for assistance and support in the laboratory. Constructive comments of past and present members of the Molecular Geo- and Palaeobiology Lab, LMU München greatly improved the project. This study was possible thanks to funding of the German Science Foundation (DFG), through the SPP 1158 “Antarktisforschung", grants Wo896/9-1,2 to G. Wörheide, and DJ1063/14-2,2 and DJ1063/17-1 to D. Janussen, respectively. MD was funded through DFG Grants DO 1742/1-1,2. SV is indebted to N. Villalobos Trigueros, M. Vargas Villalobos and S. Vargas Villalobos for their constant support during the course of the study.

